# A centrosome calcium signal is essential for mammalian cell mitosis

**DOI:** 10.1101/379792

**Authors:** Nordine Helassa, Charlotte Nugues, Robert D Burgoyne, Lee P Haynes

## Abstract

To generate a complex multicellular organism like a human requires enormous expansion in cell numbers and this is achieved predominantly through mitosis. Defects in mitosis can lead to premature ageing and cancer so understanding how it is regulated has important implications for human disease. Early data from plant and invertebrate model systems indicated that calcium (Ca^2+^) could influence mitosis. Here we explore this key question in the cell biology of mammalian cells by targeting high affinity genetically encoded Ca^2+^ sensors to mitosis specific subcellular locations. We reveal a prolonged yet spatially restricted Ca^2+^ signal at the centrosomes of mitotic cells using an actin-targeted Ca^2+^ sensor. Local depletion of Ca^2+^ at centrosomes using flash-photolysis of the caged Ca^2+^ chelator diazo-2 arrests mitosis and we provide evidence that this signal emanates from the endoplasmic reticulum. In summary, we characterize a centrosomal Ca^2+^ signal as a functionally essential input into mitosis. This extends our understanding of the complex regulatory network controlling cell division and pinpoints Ca^2+^ as an important controller of this fundamental process.

## Introduction

Mitosis is the mechanism whereby individual cells make duplicate copies of themselves and it underpins the growth, development and tissue repair of mammals (Swann, 1957). This is a fundamental cellular activity, defects in which can lead to premature ageing (Dechat, Shimi et al., 2007), aneuploidy, mitotic cell death and cancer (Potapova & Gorbsky, 2017). Much is known regarding key regulatory systems that ensure fidelity during cell division with cyclin based checkpoints (Casimiro, Crosariol et al., 2012, Lara-Gonzalez, Westhorpe et al., 2012) and mitotic specific kinases (Nigg, 2001) making up the majority of the characterized regulatory network. Ca^2+^ is a universal intracellular second messenger that is found in all plants and animals (Clapham, 2007). In most species it has a similarly ubiquitous role in controlling cellular behavior. Over thirty years ago the role of Ca^2+^ during mitosis was investigated, initially using plant cells (Hepler & Callaham, 1987, Keith, Ratan et al., 1985) and model invertebrate systems amenable to micro-manipulation and imaging techniques (Groigno & Whitaker, 1998, Poenie, Alderton et al., 1985). These studies observed both transient and global Ca^2+^ signals that appeared to correlate with specific mitotic events (nuclear envelope breakdown, metaphase-anaphase transition, cytokinesis entry). Furthermore, various functional experiments manipulating cytoplasmic Ca^2+^ during mitosis in these model systems strongly argued an essential role for Ca^2+^during normal mitotic progression (Groigno & Whitaker, 1998, Steinhardt & Alderton, 1988, Twigg, Patel et al., 1988, Wilding, Wright et al., 1996). When similar experiments were attempted in somatic mammalian cells the results were inconclusive (Hepler, 1994, Kao, Alderton et al., 1990, Poenie, Alderton et al., 1986, Ratan, Shelanski et al., 1986, Tombes & Borisy, 1989, Whitaker, 2006).

In this study we have investigated the role of Ca^2+^ during mitosis in mammalian cells by developing a refined molecular toolkit. Our approach was to take proteins implicated in specific mitotic events and tag them with the GCaMP6s Ca^2+^ sensor (Chen, Wardill et al., 2013, Helassa, Podor et al., 2016). Our aim was to spatially restrict a high affinity Ca^2+^ probe to mitosis specific locations in dividing cells which we hypothesized would permit the detection of mitosis specific Ca^2+^ signals.

## Results and discussion

Actin has a well-documented role in forming the contractile ring necessary for constriction of the plasma membrane (PM) late in mitosis during telophase as the two nascent daughter cells prepare for physical separation (Miller, 2011). Initial validation of actin-GCaMP6s demonstrated that when expressed in interphase HeLa cells it co-localised extensively with cellular actin filaments (Fig. 1*A*) and migrated at the predicted molecular weight on SDS-PAGE (Fig. expanded view 1*A*). In functional tests of actin-GCaMP6s’ ability to respond to Ca^2+^ we challenged infected cells with ionomycin/Ca^2+^ to elevate cytoplasmic Ca^2+^ chronically (Fig. 1*C* & Fig. expanded view 1*B*) or applied the Ca^2+^ mobilizing agonist histamine, which elicits well-documented Ca^2+^ oscillations in HeLa cells (Fig. 1*B* & Fig. expanded view 1*B*). In both instances we observed robust increases in actin-GCaMP6s fluorescence with histamine driving the expecting oscillatory behaviour. We also confirmed that actin-GCaMP6s expression had no deleterious effect on cell division (overall % of healthy cells completing mitosis and average time taken to transit mitosis) when compared with control, untransfected, cells (Fig. expanded view 1*C* & *D*). Remarkably, when expressed in HeLa cells, actin-GCaMP6s detected a highly localized Ca^2+^ signal during mitosis (Fig. 2). A recent study of interphase cells characterized the centrosome as having the ability to organize not only the microtubule network but also actin filaments (Farina, Gaillard et al., 2016). We therefore investigated the possibility that the probe was targeting to centrosomes in mitotic cells. Using a centrosome specific marker, mCherry-CEP135, we confirmed its co-localisation with actin-GCaMP6s during mitosis (Fig. 2*A*). This data shows both the presence of centrosome-associated actin and a region of elevated Ca^2+^ specifically at the centrosome in mitotic cells only (Fig. 2*A*). This phenomenon was observed in three cell types of widely differing lineages (HeLa, SH-SY5Y and HEK293T) suggesting that it is universal. In these analyses there are examples where an mCherry-CEP135 positive structure in a mitotic cell has no apparent corresponding actin-GCaMP6s signal (Fig. 2*A* Hela cell during mitosis). This could be due to variation in local Ca^2+^ being experienced by individual centrosomes during mitosis such that some are highly fluorescent for actin-GCaMP6s whilst others are less apparent. The Ca^2+^ detected by actin-GCaMP6s is visible from prophase and persists through to telophase (Fig. 2*B*). It therefore exhibits the unusual property of being spatially but not temporally focal. In order to prove that actin-GCaMP6s was registering a true Ca^2+^ signal at the centrosome we tested the response of cells to elevation or depression of the cytoplasmic Ca^2+^ concentration (Fig. 2*C*). These experiments proved that actin-GCaMP6s was expressed extensively in cells (as would be expected for an actin based construct) and that its fluorescence could be activated globally by ionomycin treatment in the presence of external Ca^2+^ (Fig. 2*C*). Similarly, the focal nature of the Ca^2+^ signal was confirmed by its disappearance when cells were incubated with the cell permeant Ca^2+^chelator, BAPTA-AM, which as a fast chelator would be effective at suppressing Ca^2+^ microdomains (Fig. 2*C*). Previous studies examining actin dynamics in mitotic cells (Fink, Carpi et al., 2011, Mitsushima, Aoki et al., 2010) did not observe actin specifically at centrosomes but highlighted that actin can be found in close proximity to spindle microtubules. We have shown that actin-GCaMP6s is expressed at many locations in mitotic cells (Fig. 2*C*) but detects a Ca^2+^ signal uniquely limited to the centrosome. Our observation of centrosomal actin during mitosis is new and consistent with that of centrosomal actin in interphase cells (Farina et al., 2016). We speculate that as well as nucleating actin in interphase cells, centrosomes may also exhibit this behavior during mitosis. We do not believe that the centrosome GCaMP signal is a non-specific consequence of concentrating the probe into a restricted volume for various reasons. Firstly, GCaMP6s should have effectively undetectable fluorescence at resting Ca^2+^ levels (Chen et al., 2013). Secondly, we observe actin-GCaMP6s structures in mitotic cells but they are not always present where mCherry-CEP135 is detected (Fig. 2*A* and discussion above).Thirdly, other studies using a PM targeted GCaMP did not report fluorescence in the absence of a Ca^2+^ signal (Shigetomi, Kracun et al., 2010a, Shigetomi, Kracun et al., 2010b).

**Fig. 1.**
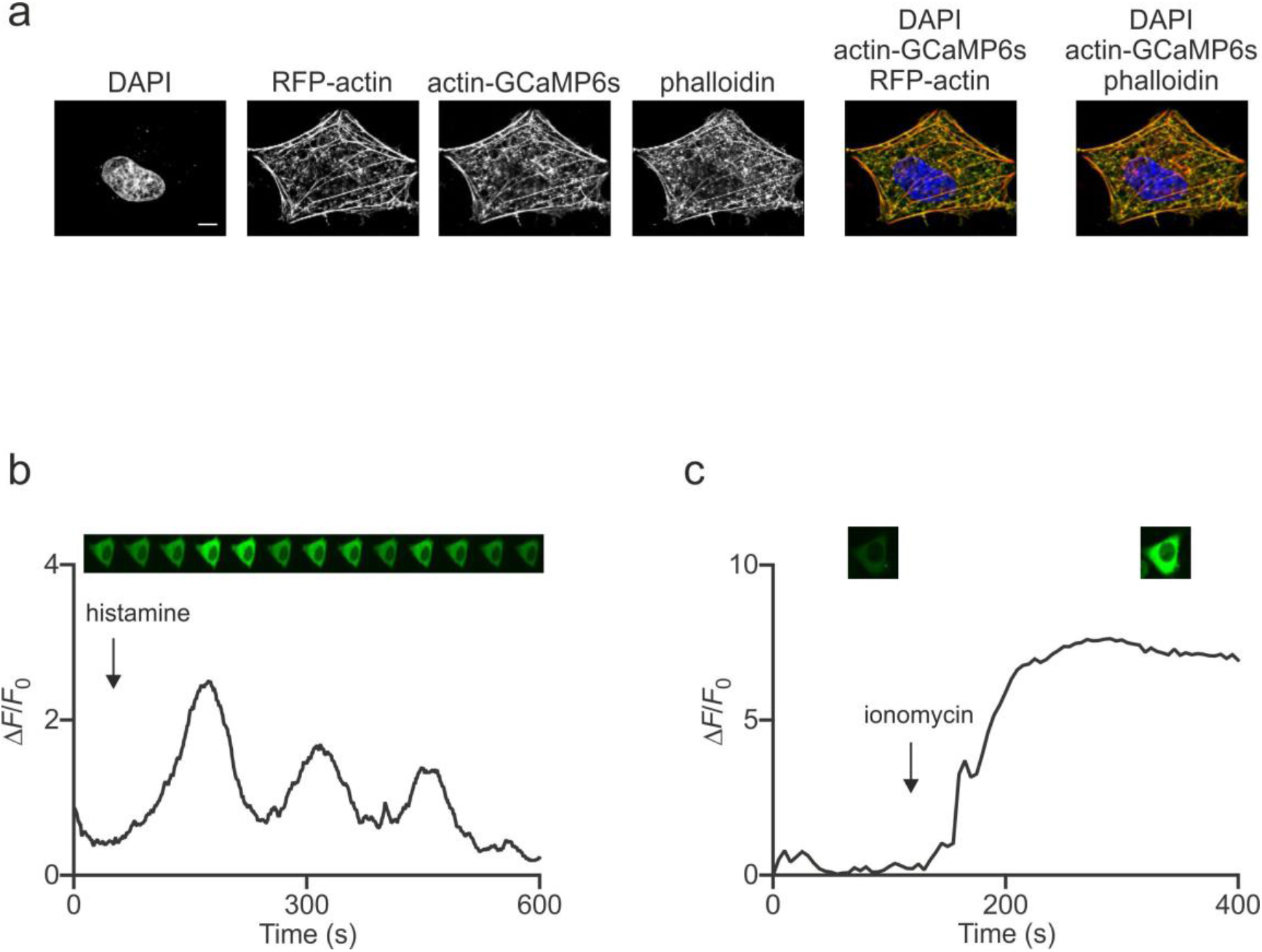
An actin-GCaMP6s Ca^2+^ sensor co-localises with actin and responds to intracellular Ca^2+^ signals. (A) Immunofluorescence confocal microscopy images of HeLa cells expressing RFP-actin and actin-GCaMP6s (DAPI, blue; RFP-actin or Alexa Fluor 647 phalloidin, red; actin-GCaMP6s, green). Scale bar, 10 μm. (*B,C*) Representative traces of HeLa cells expressing actin-GCaMP6s stimulated with (*B*) 500 μM histamine or (*C*) 10 μM ionomycin. Corresponding live-cell confocal images are presented in the top panel.

**Fig. 2.**
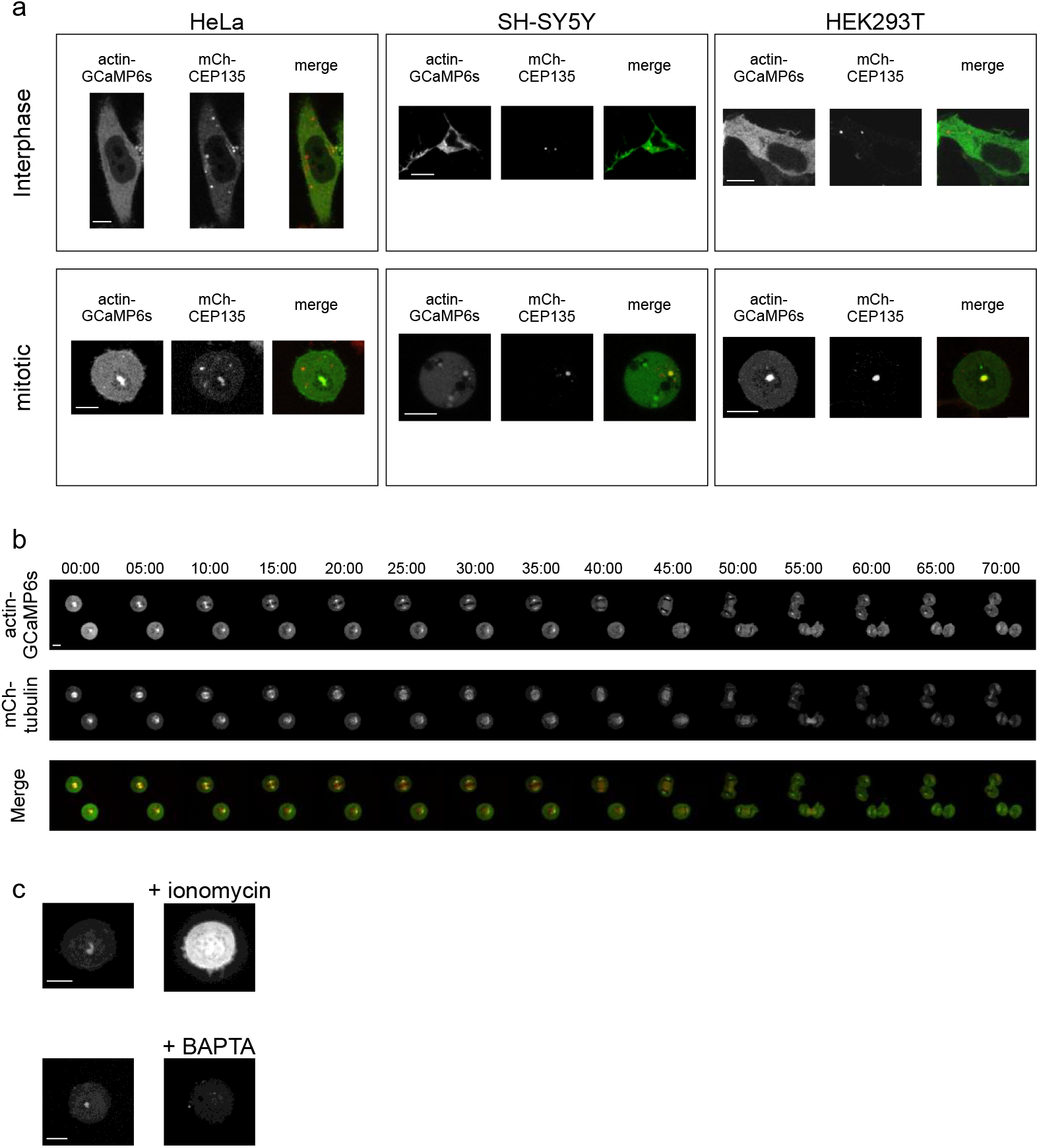
An actin-GCaMP6s probe detects a focal, persistent and specific Ca^2+^ signal at the centrosomes of mammalian cells during mitosis. (*A*) Representative confocal images of HeLa, SH-SY5Y and HEK293T cells expressing actin-GCaMP6s (green) and CEP135 centrosomal protein (red) in interphase and mitosis. Scale bar, 10 μm. (*B*) Live-cell confocal images of HeLa cells expressing actin-GCaMP6s (green) and mCherry-tubulin (red). Time stamps are displayed in minutes:seconds. (*C*) Representative confocal images of HeLa cells expressing actin-GCaMP6s challenged with 10 μM ionomycin or 10 μM BAPTA-AM. Scale bar, 10 μm.

Having characterised a reproducible centrosomal Ca^2+^ signal we next wanted to determine its functional significance. The focal nature of the Ca^2+^ meant that it would be impossible to manipulate using standard cell permeant chelators without simultaneously interfering with all potential Ca^2+^ based signaling occurring in the cell. We therefore chose to utilize uncaging of the compound diazo-2-AM (Adams, Kao et al., 1989), a cell permeant and UV photolysable Ca^2+^-chelator. This approach allowed us to activate the chelator only on one centrosome when cells were entering mitosis (Fig. 3*A-B* & Fig. expanded view 2 and Movie expanded view 1 & 2). When diazo-2-AM loaded cells were tracked following centrosomal UV irradiation only 4% of healthy (viable, non-dead) cells progressed normally through mitosis in comparison to 56% of non-irradiated diazo-2-AM loaded viable control cells from the same experiments (Fig. 3*A*). In control conditions with no diazo-2-AM loading and no UV irradiation, approximately 55% of all viable cells progressed normally through mitosis (Fig. 3*A*). This reflects the fact that in chemically synchronized HeLa cell populations a significant proportion of healthy cells fail to release from the chemical block and progress through mitosis. In these experiments, cells with no diazo-2 loading but where the uncaging UV irradiation protocol was applied to actin-GCaMP6s positive centrosomes, the number of healthy irradiated cells that progressed normally through mitosis was similar at 72% (Fig. 3*A*). UV irradiation of the centrosome in the absence of diazo-2 therefore had no deleterious influence on mitosis progression. Collectively these data show that specific uncaging of a Ca^2+^-chelator over a single centrosome in a dividing mammalian cell is sufficient to suppress the centrosomal Ca^2+^ signal and precipitate an immediate block on further progression through mitosis. The possibility exists that the specific combination of UV irradiation and the presence of diazo-2 within cells is somehow non-specifically toxic and inhibits mitosis progression however in order to focally deplete Ca^2+^ in live cells we are unable to conceive of a more suitable experimental design to the one employed here. In the original paper detailing the synthesis of diazo-2 (Adams et al., 1989), a related compound, diazo-3 was also described. Diazo-3 is the same as diazo-2 in regards to most of its chemistry (UV irradiation induced formation of reactive intermediates and release of protons) but is unable to chelate Ca^2+^. UV irradiation of diazo-3 was shown to induce only very limited covalent modification of a target substrate (a test for the formation of reactive intermediates where only 6 out of every 10^5^ lysine molecules were covalently modified) and did not exhibit pH related cellular toxicity. We are therefore confident that the effects that we have observed when using diazo-2 in this study are specifically restricted to its ability to chelate Ca^2+^. Another striking feature of these analyses is that the Ca^2+^ signal at centrosomes does not reappear following a relatively short (10’s of ms) UV uncaging protocol. Determining the precise nature of how centrosomal Ca^2+^ is generated and maintained throughout mitosis is beyond the scope of the present paper. It is nonetheless interesting to speculate about a possible mechanism. A recent study has shown that specific populations of immobile inositol 1,4,5-trisphosphate (IP3) receptors (IP_3_Rs) are responsible for active Ca^2+^ signaling at ER-PM membrane junctions (Thillaiappan, Chavda et al., 2017). It has also been reported that ER Ca^2+^ release is regulated by actin polymerization (Wang, Mattson et al., 2002). One hypothesis that we have is that centrosomal actin might perhaps be responsible for organizing IP3R signaling domains in close proximity to centrosomes in mitotic cells and that maintenance of these domains depends on continuity of the Ca^2+^ signal. Once the Ca^2+^ signal is disrupted, the connection between ER-centrosome is lost and cannot be re-established. Future work will be directed at investigating such possibilities to provide a complete mechanistic explanation of centrosome Ca^2+^.

**Fig. 3.**
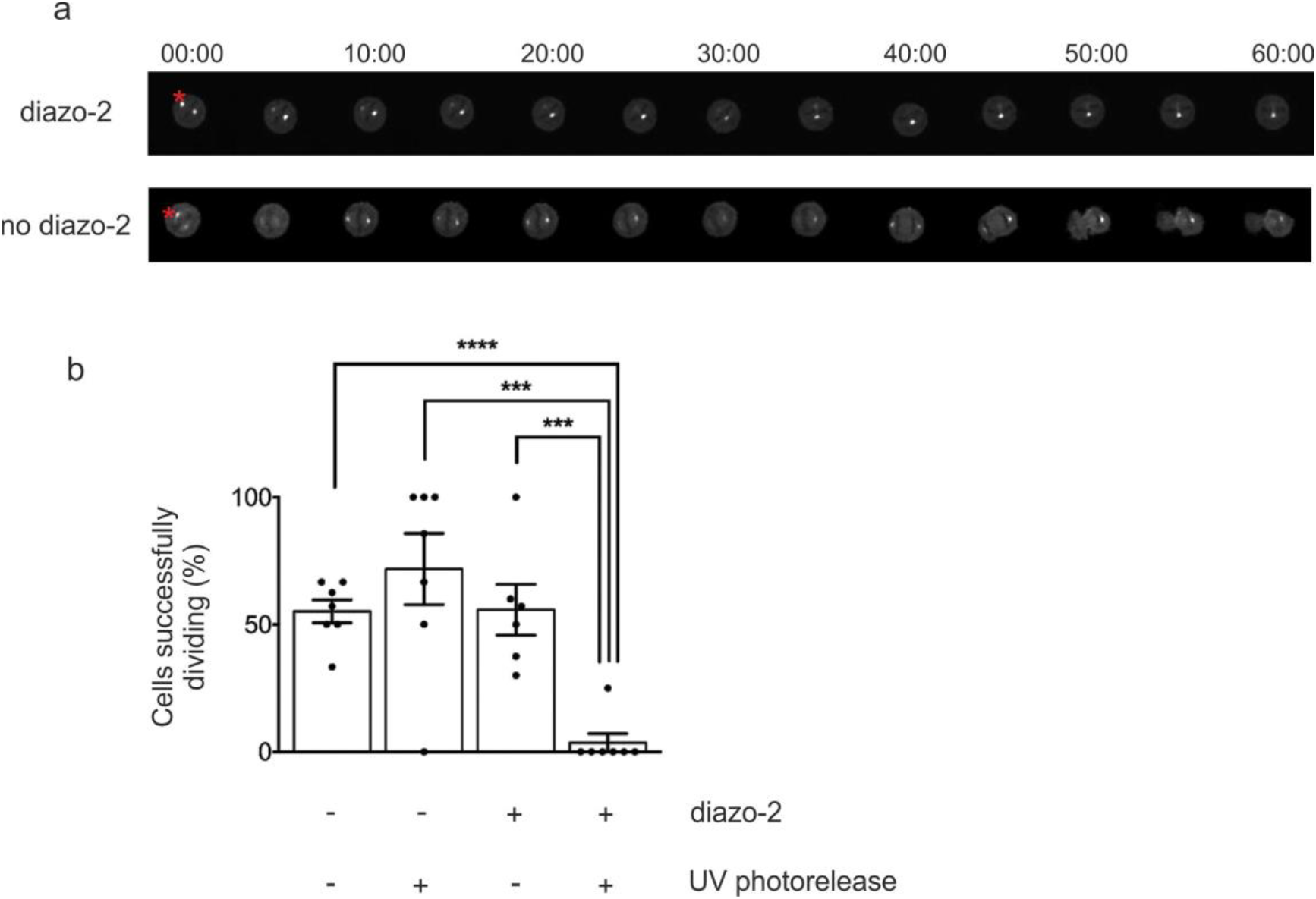
A centrosomal Ca^2+^ signal is essential for mitosis progression. (*A*) Live-cell confocal images of HeLa cells stably expressing actin-GCaMP6s. Cells were UV-irradiated at the centrosomes (red star) with or without pre-incubation with diazo-2-AM Ca^2+^chelator. Time stamps are displayed in minutes:seconds. (*B*) Quantitative analysis of the effect of Ca^2+^ depletion at the centrosomes on mitosis completion. Four different conditions were tested: HeLa cells stably expressing actin-GCaMP6s without UV-irradiation (n=39, N=7) and with UV-irradiation at the centrosomes (n=16, N=7, p<0.0001); pre-incubated with diazo-2 without UV-irradiation (n=38, N=6, p=0.0005) and UV-irradiation at the centrosomes (n=10, N=7, p=0.0003). n is the total number of cells analysed, N the number of independent experiments and p the calculated probability (p-value). Results are expressed as mean ± SEM.

Finally, we wanted to gather evidence for a likely source of Ca^2+^ that was feeding the centrosomal signal in mitotic cells. For this analysis we employed a series of standard and widely used pharmacological inhibitors of various Ca^2+^-mobilising pathways in addition to the cell permeant Ca^2+^Chelator BAPTA-AM (Fig. 4*A*, Fig expanded view 3). Many of these agents will have pleiotropic effects on cells and they cannot be used to infer a direct influence on centrosomal Ca^2+^. We therefore restrict our interpretation of the data to provide circumstantial evidence as to a likely source of cellular/extracellular Ca^2+^ that is able to influence mitosis and which could be consistent with our observations of centrosomal Ca^2+^. Application of BAPTA-AM to cells led to a significant reduction in the number of cells able to successfully complete mitosis. This result confirms the general importance of intracellular Ca^2+^ for normal completion of mitosis and is consistent with data from model cell systems (Groigno & Whitaker, 1998, Steinhardt & Alderton, 1988). Similarly, three independent methods of antagonising IP3R dependent ER Ca^2+^ release all significantly impeded completion of mitosis. The phosphatidylinositol-phospholipase C (PI-PLC) specific antagonist, ET-18-OCH3 (edelfosine)(Seewald, Olsen et al., 1990), which inhibits IP3 production elicited a significant reduction in the number of viable cells completing mitosis. This is consistent with previous work demonstrating a role for IP3 during mitosis (Becchetti & Whitaker, 1997, Groigno & Whitaker, 1998, Shearer, De Nadai et al., 1999). Similarly, treatment of cells with the IP3R inhibitor caffeine (Saleem, Tovey et al., 2014) or the sarcoplasmic reticulum Ca^2+^-ATPase pump inhibitor thapsigargin (Thastrup, Cullen et al., 1990) elicited significant impairment of mitosis completion (Fig. 4*A*, Fig. expanded view 3). Treatment of cells with the store operated Ca^2+^ entry (SOCE) inhibitor, BTP-2 (YM-58483)(Yoshino, Ishikawa et al., 2007), elicited a small, if significant, increase in cells failing to progress through mitosis. The limited effect of BTP-2 is perhaps not unexpected as SOCE entry through Orai channels is known to be down-regulated during mitosis (Smyth & Putney, 2012). The lysosomal V-ATPase inhibitor concanamycin-A which induces lysosomal Ca^2+^ depletion was without effect on mitosis. Lysosomes have recently been characterized as important Ca^2+^-signaling platforms (Galione, 2015) although our data argues against a role for lysosomal Ca^2+^ release during mitosis. We employed two cell synchronization protocols in this part of the study, double-thymidine (cells arrested in interphase) or thymidine-RO-3306 (CDK1 inhibitor which arrests cells at the G2/M boundary). The data sets are complimentary for each protocol and indicate that the Ca^2+^ signal important for mitosis therefore occurs at some point during or following prophase, that its source is the ER and that an IP3 generating agonist working through PI-PLC activation is required. These experiments were additionally performed in parallel on two independent cell lines, HeLa (Human cervical epithelial cells) and SH-SY5Y (human neuroblastoma cells). The data sets for each cell line follow identical trends, indicating that ER Ca^2+^ mobilized by IP3 is a universal requirement for mitosis progression in mammalian cells.

**Fig. 4.**
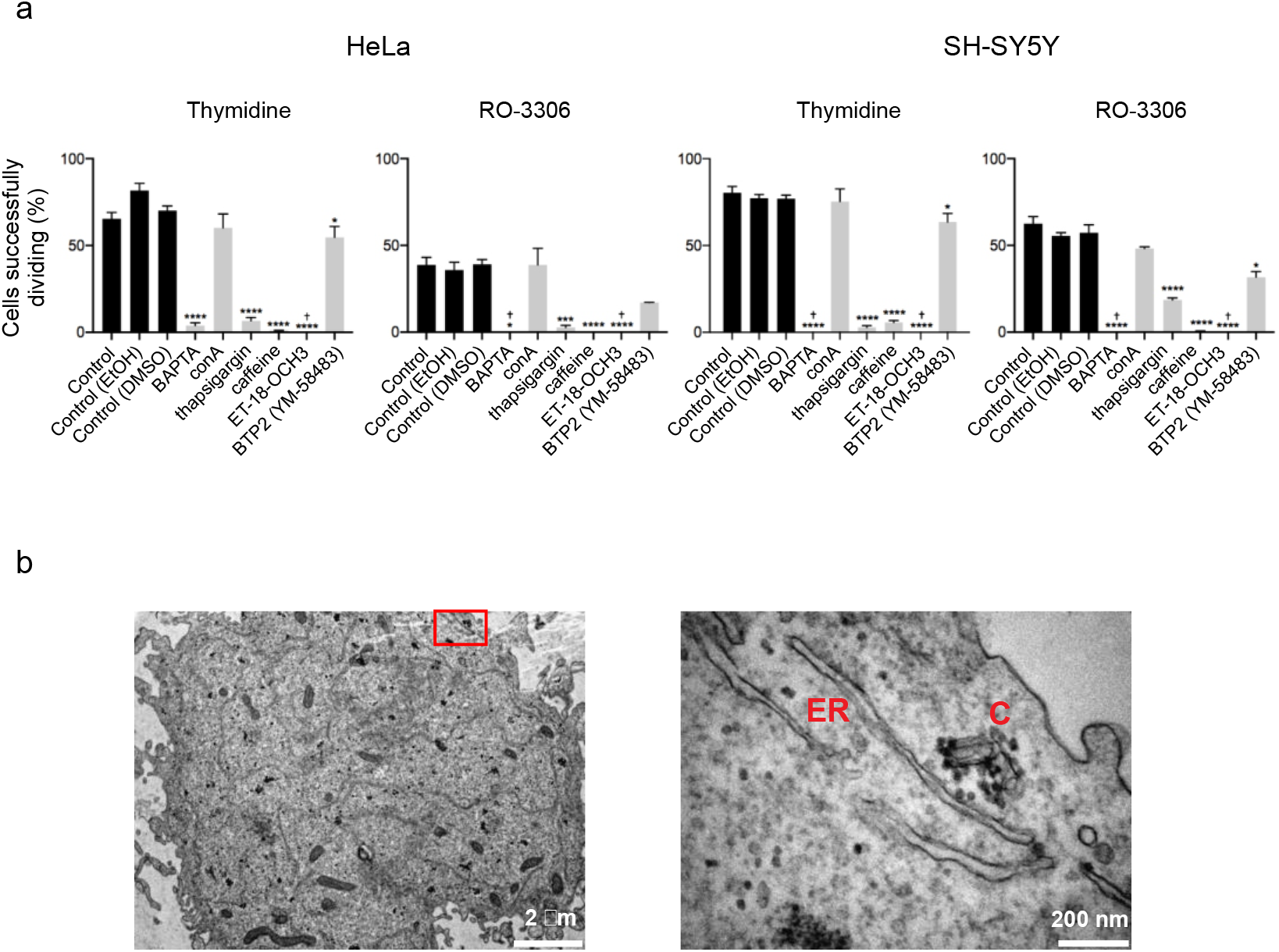
Ca^2+^ from the endoplasmic reticulum is required for mitosis progression. (*A*) Effect of pharmacological treatments targeting Ca^2+^ signaling pathways on cell division in HeLa and SH-SY5Y cells synchronized at interphase (thymidine) or G2/M (RO-3306). Results are expressed as mean ± SEM. † indicates complete cell death (*B*) Representative transmission electron microscopy image of a mitotic HeLa cell at 6K (left panel) or 60K (right panel) magnification. C=centrosome and ER=endoplasmic reticulum.

Finally, we examined the localization of centrosomes and ER in mitotic HeLa cells by transmission electron microscopy (TEM, Fig. 4*B*, Fig. expanded view 3*B*) to look for evidence that these two cellular entities might exist in close proximity, consistent with a model where the ER is the organelle driving centrosomal Ca^2+^. In plant cells (Hepler, 1980) and invertebrate model organisms (Terasaki & Jaffe, 1991) the ER has been shown to associate closely with the mitotic apparatus and spindle poles and in our studies we observed strands of ER closely apposed with centrosomes. The limited number of high quality images of mitotic cells that we were able to generate for this analysis restricted us to a purely qualitative examination of the TEM data. This did however reveal that ER was the most frequently observed, morphologically distinguishable, cellular organelle located close to centrosomes, consistent with the pharmacological treatment data and our assertions regarding the source of centrosomal Ca^2+^

In this study we have employed spatial restriction of a genetically encoded Ca^2+^ sensor to detect a focally restricted centrosome based Ca^2+^ signal in mitotic cells. This work connects the cell biology of Ca^2+^ signaling with mitosis in mammalian cells and has important implications for furthering our understanding of normal growth, development and ageing. This is likely a universal phenomenon and, crucially, is functionally essential during mitosis. Centrosomal Ca^2+^ represents an entirely novel control mechanism which opens new avenues of investigation into modulating mitosis.

## Materials and Methods

### Plasmids

pTagRFP-actin was purchased from Evrogen. pcDNA3.1(+) was purchased from Invitrogen. pGP-CMV-GCaMP6s was a gift from Douglas Kim (Addgene plasmid # 40753). pHIV-dTomato was a gift from Bryan Welm (Addgene plasmid # 21374). pMDLg/pRRE and pRSV-Rev were gifts from Didier Trono (Addgene plasmid # 12251 and # 12253). pCMV-VSV-G was a gift from Bob Weinberg (Addgene plasmid # 8454). pANT7_cGST-CEP135 was purchased from DNASU (# HsCD00640042)(Seiler, Park et al., 2014).

### Generation of pcDNA3.1(+)-actin-GCaMP6s and pmCherry-CEP135

To create pcDNA3.1(+)-actin-GCaMP6s, we first amplified the human cytoplasmic β-actin gene by PCR from pTagRFP-actin using NEB Phusion polymerase (Forward, 5’- ATATAAGCTTACCATGGATGATGATATCGCCG-3’; Reverse, 5’- ATGGATCCGAAGCATTTGCGGTGGA-3’) and cloned it into pcDNA3.1(+) by restriction-ligation using HindIII/BamHI and T4 DNA ligase (NEB). Then, the gcamp6s gene was amplified by PCR from pGP-CMV-GCaMP6s using Phusion polymerase (Forward, 5’-ATATGCGGCCGCATGACTGGTGGACAGCAAATG-3’; Reverse, 5’-ATATGGGCCCTCACTTCGCTGTCATCATTTGTAC-3’) and cloned into pcDNA3.1(+)-actin by restriction-ligation using NotI/ApaI and T4 DNA ligase. Integrity and localisation of actin-GCaMP6s in mammalian cells was verified by Western blot and immunofluorescence (see below). To create pmCherry-CEP135, we first amplified the cep135 gene by PCR from pANT7_cGST-CEP135 using Phusion polymerase (Forward, 5’-GGGCCCAATGACTACAGCTGTAGAGAG-3’; Reverse, 5’-GGATCCCTACACATTTCTATGTTCAGGAG-3’) and cloned into pmCherry-C1 by restriction-ligation using ApaI/BamHI and T4 DNA ligase. All molecular constructs were verified by DNA sequencing (The Sequencing Service, University of Dundee, UK).

### Cell culture and synchronisation

Cells were cultured in DMEM (HeLa and HEK293T) or DMEM/F12 (SH-SY5Y) containing 10% (v/v) Heat inactivated FBS (Life Technologies), 1% (v/v) non-essential amino-acids (Life Technologies), penicillin/streptomycin (100 U/ml, 100 μg/ml, respectively), at 37 °C in an atmosphere of 5% CO2. Cells were synchronised with 2 mM thymidine for 20-24 hours, released with 25 μM deoxycytidine for 5-6 hours (transfection with Lipofectamine 2000 was performed during this step when necessary, following the manufacturer’s recommendations) and blocked with either 2 mM thymidine (interphase block), 10 μM RO-3306 (G2/M block) or 35 μg/ml nocodazole (pro-metaphase block) for 15-24 hours. Cells were washed 5 times with culture medium to release the block before confocal imaging.

### Validation of actin-GCaMP6s as an actin targeted calcium sensor

#### Immunofluorescence staining

Cells co-transfected with pcDNA3.1(+)-actin-GCaMP6s and pTagRFP-actin were fixed with 4% (w/v) formaldehyde (in phosphate-buffered saline, PBS) for 15 minutes, washed with PBS and permeablised with 1% (w/v) Bovine Serum Albumin (BSA), 0.1% (v/v) Triton X-100 (in PBS) for 5 minutes. Cells were washed with PBS, blocked in 5% (w/v) BSA for 10 minutes and immunostained using mouse monoclonal anti-GFP antibody (Roche) and Alexa Fluor 647 Phalloidin (Life Technologies) for 1 hour. Cells were washed with PBS and treated with AlexaFluor 488 goat anti-mouse secondary antibody (Life Technologies) for 1 hour. Cells were washed with PBS and coverslips were mounted on slides using Prolong Gold DAPI anti-fade glycerol (Life Technologies). Microscopic observation was made on a Zeiss LSM 800 with Airyscan confocal microscope equipped with a Zeiss AxioObserver Z1, a 63x/1.4 Plan-Apochromat oil immersion objective and diode lasers as excitation light source (405 nm for DAPI; 488 nm for actin-GCaMP6s; 561 nm for RFP-actin; 640 nm for Phalloidin). Emitted light was collected through Variable Secondary Dichroics (VSDs) onto a GaAsP-PMT detector. Images were acquired using Zen Blue software and analysed on ImageJ after Airyscan processing.

#### Western blot analysis

HeLa cells transfected with GCaMP6s or actin-GCaMP6s were lysed using RIPA buffer. Proteins from the lysates were separated on SDS-PAGE (NuPAGE 4–12% (w/v) Bis-Tris, NuPAGE MOPS SDS running buffer, Life Technologies) and electrophoretically transferred onto LI-COR Odyssey nitrocellulose membranes. Membranes were blocked with 5% (w/v) non-fat milk in Tris-buffered saline/0.2% (v/v) Tween-20 (TBST) for 30 minutes and incubated with mouse monoclonal anti-GFP primary antibody (1:1000, Roche mouse anti-GFP, Cat# 11814460001, clones 7.1 and 13.1, lot 10521400) for 1 hour. Excess primary antibody was removed by TBST washes before incubation with peroxidase-conjugated anti-mouse IgG secondary antibody (1:1000, Sigma) for 1 h at room temperature. Excess secondary antibody was removed by TBST washes and antibody binding was detected using Pierce ECL western blotting substrate (Thermo Scientific) on a ChemiDoc XRS+ (Biorad).

#### Calcium imaging

HeLa cells stably expressing actin-GCaMP6s were plated on 35 mm glass-bottom dishes and challenged with 10 μM ionomycin or 500 μM histamine. Fluorescence over time was recorded on a 3i Marianas spinning-disk confocal microscope equipped with a Zeiss AxioObserver Z1, a 40x/1.3 Plan-Apochromat oil immersion objective and a 3i Laserstack as excitation light source (488 nm, for actin-GCaMP6s). Emitted light was collected through a single bandpass filter (Yokogawa CSU-X filter wheel) onto a CMOS camera (Hamamatsu, ORCA Flash 4.0). Images were collected at 1 frame every second (histamine) or every 5 seconds (ionomycin) using SlideBook 6 software and processed on ImageJ. Data obtained from 10 cells was normalised and plotted on GraphPad Prism 6.

### Generation of stable HeLa cells expressing actin-GCaMP6s

Actin-GCaMP6s gene was amplified by PCR from pcDNA3.1(+)-actin-GCaMP6s using Phusion polymerase (Forward, 5’-CATCATCTAGAGCTGGCTAGCGTTTAAAC-3’; Reverse, 5’- GTAGTAACGTTCTG ATCAGCGGGTTTAAAC-3’) and cloned into pHIV-dTomato by restriction-ligation using XbaI/(AclI or ClaI) and T4 DNA ligase. During the process, dTomato is replaced by actin-GCaMP6s. All molecular constructs were verified by DNA sequencing (The Sequencing Service, University of Dundee, UK).

To generate stable HeLa cells expressing actin-GCaMP6s, we used a 3^rd^ generation lentivirus system: pHIV-actin-GCaMP6s (20 μg) was co-transfected with pMDLg/pRRE (10 μg), pRSV-Rev (5 μg) and pCMV-VSV-G (6 μg) using Lipofectamine 2000 following the manufacturer’s recommendations in HEK293T cells (10 cm tissue culture dish). Supernatant containing the lentivirus particles was collected 48 hours post-transfection, centrifuged and clarified by syringe filtration (0.45 μm). HeLa cells were infected with actin-GCaMP6s lentivirus in a 6-well plate for 24 hours. The stable cell line was then expanded and used for experiments.

### Localisation of actin-GCaMP6s and centrosomes using live-cell confocal imaging

HeLa, SH-SY5Y and HEK293T cells were plated on 35-mm glass bottom culture dishes and synchronised using double thymidine (interphase) or thymidine-nocodazole (mitotic) blocks as described above. During the first release, cells were co-transfected with actin-GCaMP6s and mCherry-CEP135 using Lipofectamine 2000 following manufacturer’s recommendations. Cells were imaged on a Zeiss LSM 800 laser scanning confocal microscope equipped with a Zeiss AxioObserver Z1, a 63x/1.4 Plan-Apochromat oil immersion objective and diode lasers as excitation light source (488 nm, for actin-GCaMP6s; 561 nm, mCherry-CEP135). Emitted light was collected through Variable Secondary Dichroics (VSDs) onto a GaAsP-PMT detector. Images were acquired using Zen Blue software (Zeiss) and processed on ImageJ.

### Laser flash photolysis of diazo-2 (caged calcium chelator) in live-cell confocal imaging

HeLa cells stably expressing actin-GCaMP6s were plated on 35 mm glass-bottom dishes and synchronised using thymidine-nocodazole protocol. One hour before the second release, cells were loaded with 2 μM diazo-2-AM (Molecular Probes, a kind gift from Prof Alexei Tepikin, University of Liverpool, UK) for at least 30 minutes at 37 °C.

Cells were examined in a 3i Marianas spinning-disk confocal microscope equipped with a Zeiss AxioObserver Z1, a 40x/1.3 Plan-Apochromat oil immersion objective and a 3i Laserstack as excitation light source (405 nm, for diazo-2 photolysis; 488 nm, for actin-GCaMP6s). Emitted light was collected through single bandpass filters (Yokogawa CSU-X filter wheel) onto a CMOS camera (Hamamatsu, ORCA Flash 4.0). Experiments were carried out at 37 °C and 5% (v/v) CO2 (OKO lab incubation chamber).

Photolysis of diazo-2 was performed by illumination with 405 nm laser light at 10% power for 10 ms for rapid chelation of [Ca^2+^] at the centrosome (elliptical region of Interest, ROI) after 1 to 3 frames. Then, cell division was monitored by collecting images every minute using SlideBook 6 software and processed on ImageJ and GraphPad Prism 6.

### Pharmacological treatments in live-cell confocal imaging

HeLa and SH-SY5Y cells were plated onto 24-well tissue culture plate and synchronised with double thymidine or thymdine-RO-3306 protocols as described above. After the second release, cells were treated with 25 μM BAPTA-AM, 2 μM thapsigargin, 25 nM concanomycin-A, 10 mM caffeine, 100 μM ET-18-OCH3, 10 μM YM-58483 or vehicle controls. Cells were imaged on a 3i Marianas spinning-disk confocal microscope equipped with a Zeiss AxioObserver Z1 and a 10x/0.45 Plan-Apochromat objective. Experiments were carried out at 37 °C and 5% (v/v) CO2 (OKO lab incubation chamber). Cell division was monitored by collecting transmitted light images onto a CMOS camera (Hamamatsu, ORCA Flash 4.0) every 5 minutes using SlideBook 6 software and processed on ImageJ.

### Transmission Electron Microscopy

Samples were prepared for transmission electron microscopy (TEM) as follows. HeLa cells were plated on a 10 cm dish and synchronised using thymidine-nocodazole protocol as described above. Mitotic cells (“shake-off” method) were fixed in 2.5% (w/v) glutaraldehyde in 0.1 M phosphate buffer pH 7.4 (PB) in Pelco Biowave (Ted Pella Inc.) 1 minute “on”, 1 minute “off”, 1 minute “on”, 100 W, 20 Hg. Cells were then washed twice in 0.1 M PB before embedding in 3% (w/v) agarose. Agarose embedded cell pellets were cut into small cubes before being post fixed and stained with reduced osmium, (2% (w/v) osmium tetroxide in dH2O + 1.5% (w/v) Potassium Ferrocyanide in 0.1 M PB) in Biowave 20 seconds “on”, 20 seconds “off”, 20 seconds “on”, 20 seconds “off”, 20 seconds “on”, 100 W, 20 Hg. This was followed by a second osmication step (2% (w/v) in ddH2O, 20 seconds “on”, 20 seconds “off”, 20 seconds “on”, 20 seconds “off”, 20 seconds “on”, 100 W, 20 Hg). Samples were incubated overnight at 4 °C in aqueous 1% (w/v) uranyl acetate. To prevent precipitation artifacts, the cells were washed for a minimum of 5 x 3 minutes with ddH2O between each of the staining steps described. The next day cells were washed in ddH2O before dehydrated in a graded ethanol series of 30%, 50%, 70%, 90% (v/v) in ddH2O for 5 minutes each, followed by 2 x 5 minutes 100% ethanol. Samples were then infiltrated with TAAB medium Premix resin at ratios of 1:1 with resin:100% ethanol for 30 minutes and finally samples were incubated in 100% resin for 2 x 30 minutes, before embedding the pellets in fresh 100% resin in silicone moulds and Beem capsules. Samples were cured for 48 hours at 60 °C.

Ultrathin serial section (70-75nm) were cut on an UC6 ultramicrotome (Leica, Vienna) collected on formvar coated copper grids, before viewing at 120 KV in a FEI Tecnai G^2^ Spirit. Images were taken using a MegaView III camera using analysis software at various magnifications. Multiple Image Alignment (MIA) was used on some images to create a high-resolution overview of areas of interest.

### Data analysis and statistics

Results are expressed as mean ± SEM unless indicated otherwise. All experiments were performed at least in triplicates. Number of experiments and total number of cells analysed are given in the figure legends. Significance level was obtained using an unpaired two-tailed Student’s *t*-test (GraphPad Prism 6 software) and p values in the figures are represented by stars (*p < 0.05, **p < 0.01, ***p < 0.001, ****p < 0.0001, ns for non-significant).

## Acknowledgements

This work was funded by Leverhulme Trust project grant RPG-2014-194 awarded to L.P.H and a Wellcome Trust Prize PhD studentship awarded to C.N. We would like to thank Prof Alan Morgan and Prof Alexei Tepikin (Department of Cellular and Molecular Physiology, University of Liverpool) for their insightful comments on the manuscript. All light and electron microscopy was performed at The Biomedical Imaging Facility, Institute of Translational Medicine, University of Liverpool. We thank Dr Alison Beckett for her assistance with TEM work at the EM Unit.

## Author contributions

N.H. performed all experimental work, analysed the data and wrote the manuscript. C.N. analysed the data and wrote the manuscript. R.D.B designed the study and wrote the manuscript. L.P.H designed the study, analysed the data and wrote the manuscript.

## Expanded view material

**Fig. expanded view 1.**
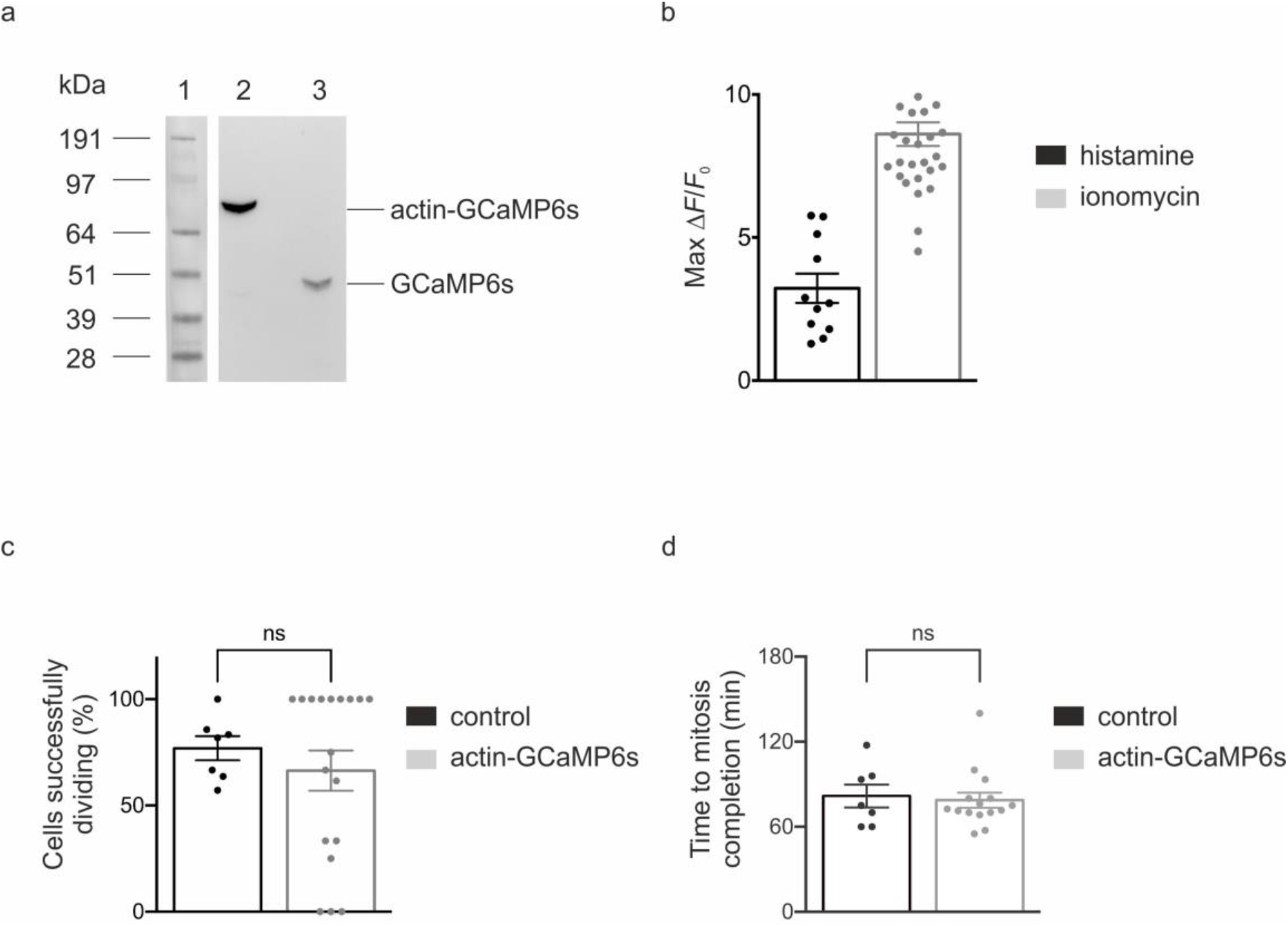
Characterisation of actin-GCaMP6s expression, functionality and cytotoxicity in HeLa cells. (*A*) HeLa cells expressing actin-GCaMP6s or GCAMP6s alone were processed for analysis by SDS-PAGE and Western blotting with anti-GFP antibody. Actin-GCaMP6s migrated at the predicted molecular weight of 91 kDa. (*B*) Peak actin-GCaMP6s fluorescence signal observed in cells treated with 500 μM histamine (n=11, N=3) or 10 μM ionomycin in the presence of 1.8 mM external Ca^2+^ (n=30, N=4). (*C*) HeLa cells expressing actin-GCaMP6s were synchronized with thymidine-nocodazole and released from cell cycle arrest. The number of cells successfully completing mitosis (defined as all healthy cells that entered mitosis and successfully completed cytokinesis) were scored as a % of the total number of healthy cells analysed (n=60, N=18) and compared to control non-transfected cells (n=46, N=7, p=0.5107). In all analyses there was a proportion of visible cell death likely due to the chemical block treatment. Non-viable cells (ignored from all cell counts) were identified as cells that failed to enter/complete mitosis (see above definition of healthy cells) in combination with having one or both of the following morphological features in brightfield images: cellular condensation, membrane rupture. (*D*) Data from (C) was additionally analysed to determine mitosis transit time (time from entry into Prophase to exit from cytokinesis) in control versus actin-GCaMP6s expressing cells. Delays in mitosis completion are often indicative of compromised cell viability. We observed no statistically significant difference in the average time taken for control or actin-GCaMP6s expressing cells to enter mitosis and exit cytokinesis. n is the total number of cells analysed, N the number of independent experiments performed and p the calculated probability (p-value). Results are expressed as mean ± SEM.

**Fig. expanded view 2.**
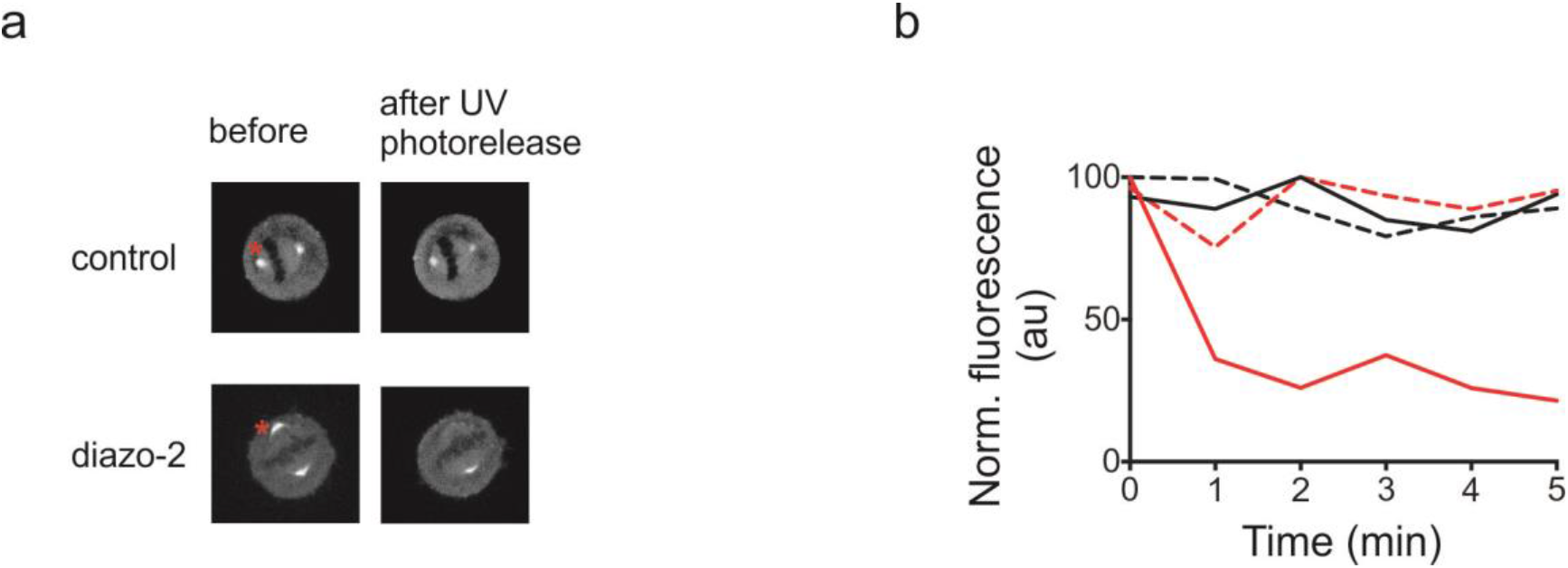
Flash photolysis of centrosomes during mitosis leads to a specific depletion of actin-GCaMP fluorescence only in diazo-2 loaded cells. (A) HeLa cells stably expressing actin-GCaMP6s were synchronized with thymidine-nocodazole and incubated with diazo-2 or left untreated (control). Cells were imaged at metaphase and a single centrosome UV irradiated (red asterisk). (*B*) Actin-GCaMP fluorescence traces corresponding to cells from (*A*). Fluorescence signal following UV irradiation (time =0) at the irradiated (solid black line) and non-irradiated (dashed black line) centrosomes of the control cell from (*A*). Fluorescence signal at the irradiated (solid red line) and non-irradiated (dashed red line) centrosomes of the diazo-2 loaded cell from (A).

**Fig. expanded view 3.**
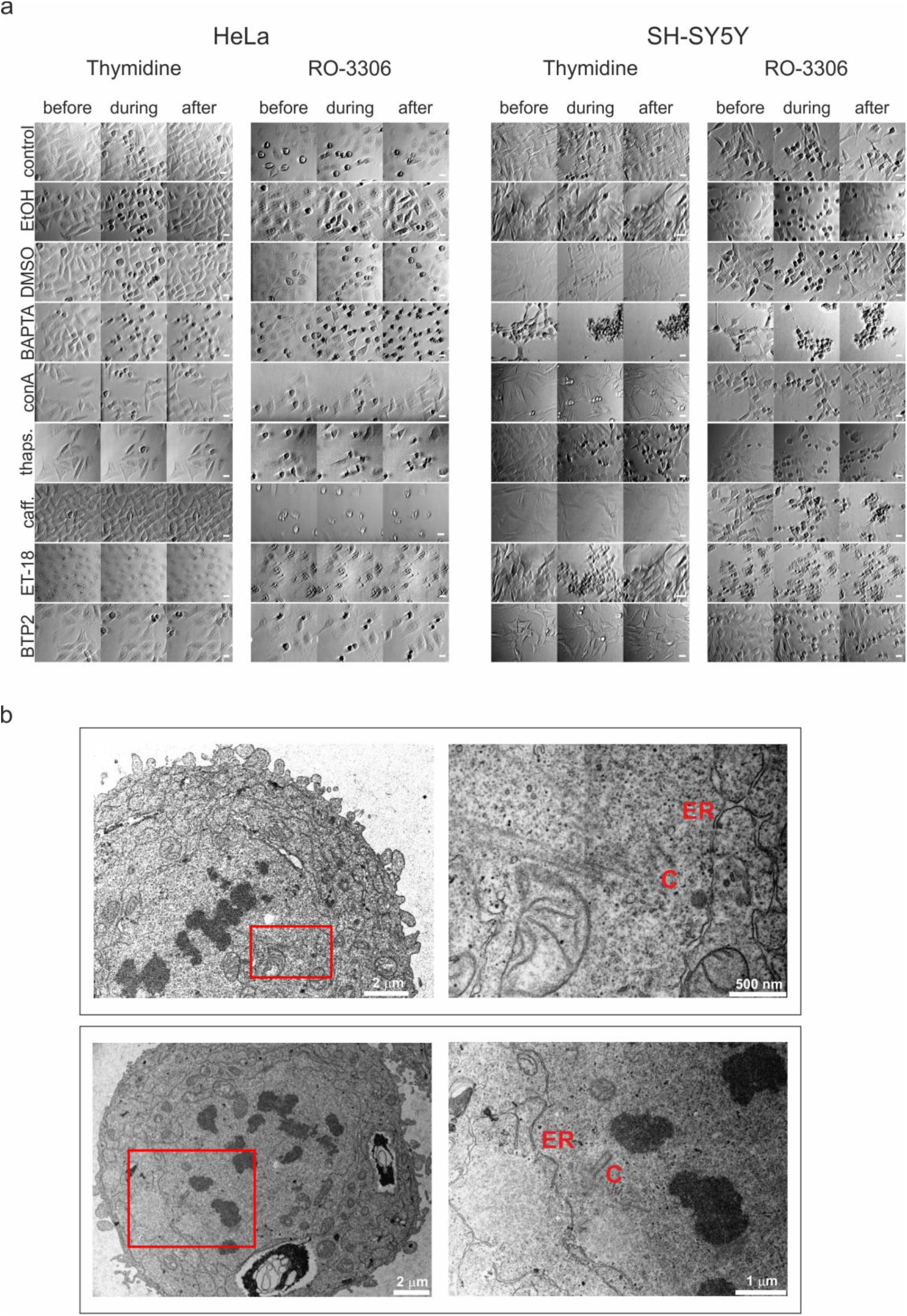
The endoplasmic reticulum is observed in close proximity to centrosomes in mitotic HeLa cells and pharmacological agents blocking ER Ca^2+^ signals induce mitotic arrest. (*A*) Brightfield images (before, during and after mitosis) of HeLa and SH-SY5Y cells synchronized with double thymidine or thymidine/RO-3306 and treated with various agents intended to disrupt specific cellular Ca^2+^ signaling pathways or vehicle controls (EtOH and DMSO). The cell counting results depicted in Fig. 4*A* histograms were derived from these data. (*B*) Examples of transmission electron micrographs taken from sections of mitotic HeLa cells. Endoplasmic reticulum (“ER”) was often observed in close proximity to centrosomes (“C”) in these samples. The higher magnification images shown in the right hand panel correspond to the regions of interest defined by red rectangles shown in the lower magnification pictures of the left hand panel.

**Movie expanded view 1. Flash photolysis of Diazo-2 at centrosomes blocks mitosis.** HeLa cells stably expressing actin-GCaMP6s were synchronized using thymidine-nocodazole and incubated with diazo-2. Cells were imaged at metaphase and a single centrosome UV irradiated (red asterisk).

**Movie expanded view 2. UV-irradiation at centrosomes does not affect cell division.** HeLa cells stably expressing actin-GCaMP6s were synchronized using thymidine-nocodazole. Cells were imaged at metaphase and a single centrosome UV irradiated (red asterisk).

